# Disrupting different *Distal-less* exons leads to ectopic and missing eyespots accurately modeled by reaction-diffusion mechanisms

**DOI:** 10.1101/183491

**Authors:** Heidi Connahs, Sham Tlili, Jelle van Creij, Tricia Y. J. Loo, Tirtha Banerjee, Timothy E. Saunders, Antónia Monteiro

## Abstract

Eyespots on the wings of nymphalid butterflies represent colorful examples of the process of pattern formation, yet the developmental origins and the mechanisms behind eyespot differentiation are still not fully understood. Here we re-examine the function of *Distal-less (Dll)* in eyespot development, which is still unclear. We show that CRISPR-Cas9 induced exon 2 mutations in *Bicyclus anynana* leads to exon skipping and ectopic eyespots on the wing. Exon 3 mutations, however, lead to null/missense transcripts, missing eyespots, lighter wing coloration, loss of scales, and a variety of other phenotypes implicating *Dll* in the process of eyespot differentiation. Reaction-diffusion modeling enabled exploration of the function of *Dll* in eyespot formation, and accurately replicated a wide-range of mutant phenotypes. These results confirm that *Dll* is a required activator of eyespot development, scale growth and melanization and point to a new mechanism of alternative splicing to achieve *Dll* over-expression phenotypes.

---

The genetic and developmental origins of the bullseye color patterns on the wings of nymphalid butterflies are still poorly understood. Eyespots originated once in ancestors of this butterfly lineage, around 90 million years ago ^1-3^, to most likely function as targets for deflecting predators away from the butterfly’s vulnerable body ^1,4,5^. Eyespots may have originated via the co-option of a network of pre-wired genes because several of the genes associated with eyespots gained their novel expression domain concurrently with the origin of eyespots ^3^. Some of these genes have since lost their expression in eyespots, without affecting eyespot development, suggesting that they did not play a functional role in eyespot development from the very beginning^3^. Yet, one of the genes, ***Distal-less (Dll)***, has remained associated with eyespots in most nymphalid species examined so far, suggesting that it may have played a functional role in eyespot origins^3, 6^.

The function of *Dll* in eyespot development was initially investigated in *B. anynana* using transgenic over-expression, RNAi, and ectopic expression tools ^7^. Overexpressing *Dll* in *B. anynana* led to the appearance of small additional eyespots on the wing as well as larger eyespots, whereas *Dll* down-regulation produced smaller eyespots, strongly implicating *Dll* as an activator of eyespot development ^7^. However, a recent study using CRISPR-Cas9 to knock-out *Dll* function in the painted lady butterfly, *Vanessa cardui* contradicted these findings. Zhang and Reed (2016)^8^ found that using two guides to disrupt exon 2 in *Dll* led to the appearance not only of distally extended eyespots but also of ectopic eyespots developing in novel locations on the wing. These observations led to a conclusion that *Dll* represses eyespot development. In addition, these researchers also showed that targeting the same exon in another butterfly, *Junonia coenia*, produced darker wing pigmentation, whereas the exact same phenotype was obtained via ectopic expression of *Dll* in the wings of *B. anynana* ^7^ and in the wings of *J. orythia*, a close relative of *J. coenia* ^9^. One possibility for the discrepancies seen across species is that *Dll* has precisely opposite functions in the different butterfly species. Another possibility, which we believed more likely, is that the outcomes of genome editing may depend on the particular site that is targeted in the genome to disrupt the gene’s function.

In order to clarify the function of *Dll* in *B. anynana*, we separately targeted both exon 2 (using single guide RNAs Sg1 and Sg2) and exon 3 (using Sg3), within the homeobox (Fig. 1a). While screening potential mutants we paid special attention to areas where *Dll* expression was previously detected in *B. anynana*. These areas included the antennae, thoracic, and abdominal legs ^10,11^ eyespot centers ^12,13^ eyespot black discs ^13,14^ and the wing margin^12^ (Fig. 1b,c,d). We predicted that targeting different exons in *B. anynana* would lead to different phenotypes.

**Figure 1.**
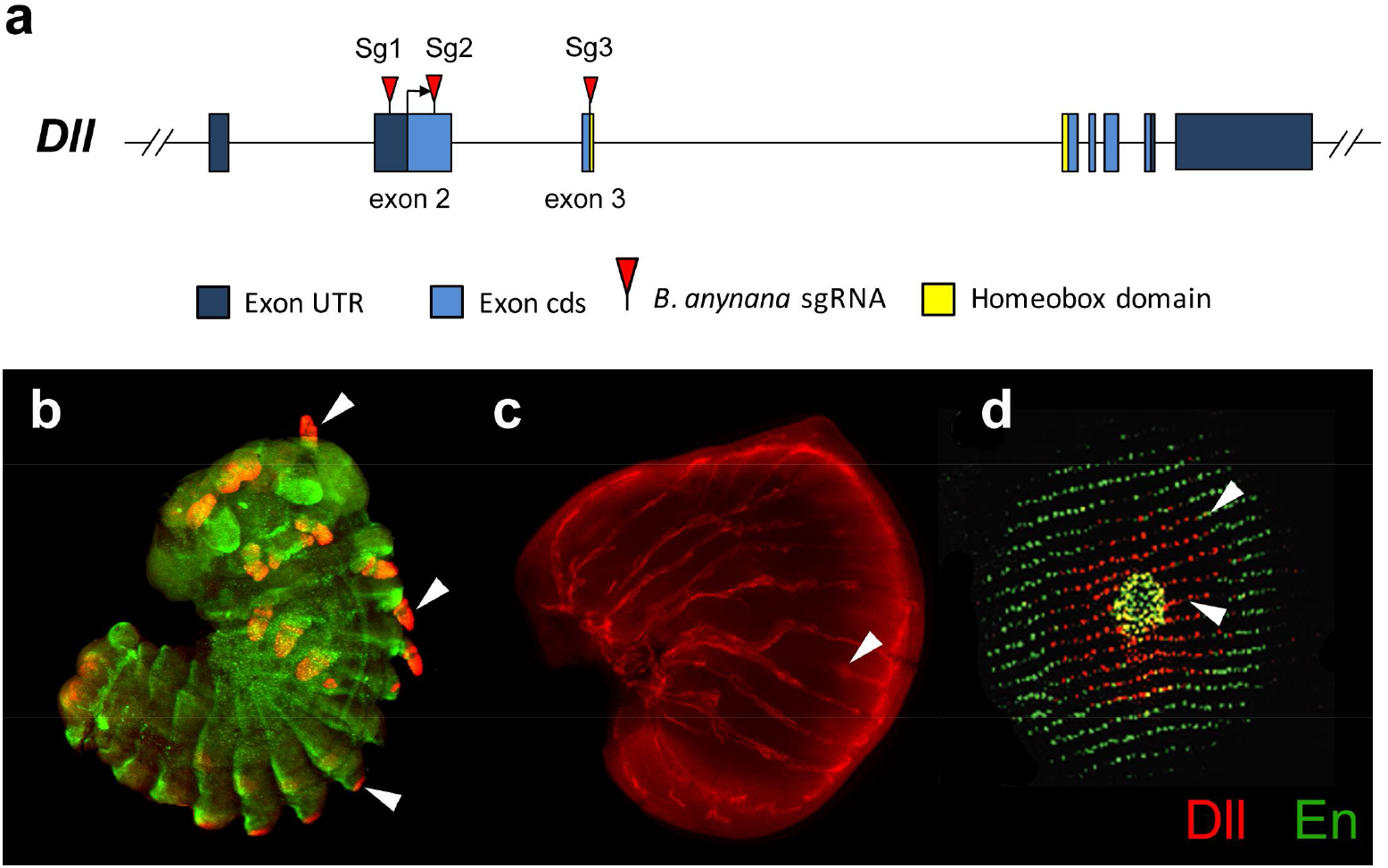
Expression of Distal-less in embryos, larval and pupal wings. **(a)** *Dll* gene structure indicating the exons targeted by guide-RNAs in this work (red triangles). **(b**) Dll (red) is expressed in antennae, thoracic legs, and abdominal prolegs of embryos (arrowheads). Engrailed (En, green) is also expressed in embryos. **(c)** Dll is expressed in eyespot centers (arrowhead) and along the wing margin in late larval wings. **(d)** Dll is expressed in eyespot centers (arrowhead) and in black scale cells of pupal wings. En is expressed in the eyespot center and area of the gold ring.

We complemented this approach with theoretical analysis of eyespot patterning. Reaction-diffusion approaches have successfully been used to model the differentiation of eyespot centers^15,16^ however, the components of these models have not been mapped to specific molecules nor have the models been tested under controlled experimental perturbation, e.g. by altering the local distribution of some of the required components. Our reaction-diffusion modeling enabled us to integrate the spatial information from morphogenetic inputs and the role of *Dll* in eyespot shape and positioning.

## Results

To confirm guide RNA efficiency *in vitro* we purified genomic amplicons of *Dll*, containing either exon 2 or exon 3, and treated them with the respective guide RNAs and with Cas9 protein. The resulting products, when run on a gel, showed two bands of the predicted sizes for Sg1 and Sg3 and a faint band for Sg2 (Supplementary Fig. 1) confirming that the CRISPR-Cas9 system was introducing double-strand breaks in the targeted sequences.

### *Dll* exon 3 mutants produced loss of function phenotypes

Embryonic injections of Sg3 targeting the *Dll* homeobox sequence on exon 3 (Fig. 1a; Table 1) led to a variety of adult phenotypes (Fig. 2, Table 2). The most striking mutants displayed complete loss of eyespots (Fig. 2a,b) followed by eyespots with significant developmental perturbations. Altered or lighter scale pigmentation, associated with the eyespot mutations, appeared to correspond to the extent of the mutant clones. Depending on their location, the lighter patches of wing tissue (i.e., the presumptive *Dll* null clones) had remarkable effects on pattern formation. Eyespots vanished when mutant patches covered the location of the eyespot centers (Fig. 2a,b), and mutant patches led to split eyespots with mutant tissue bisecting the two eyespot centers (Fig. 2c). Some patches also had lighter grey-blue scale pigmentation (Fig. 2d), lacked cover scales, or both cover and ground scales (Fig. 2e). In addition to wing mutations we observed appendage defects that would be expected from a *Dll* knockout^17,18^. A number of mutants exhibited reduced to barely noticeable stumps, legs with missing tarsi (Supplementary Fig. 2a) and deformed antennae with missing tips (Supplementary Fig. 2b).

**Figure 2.**
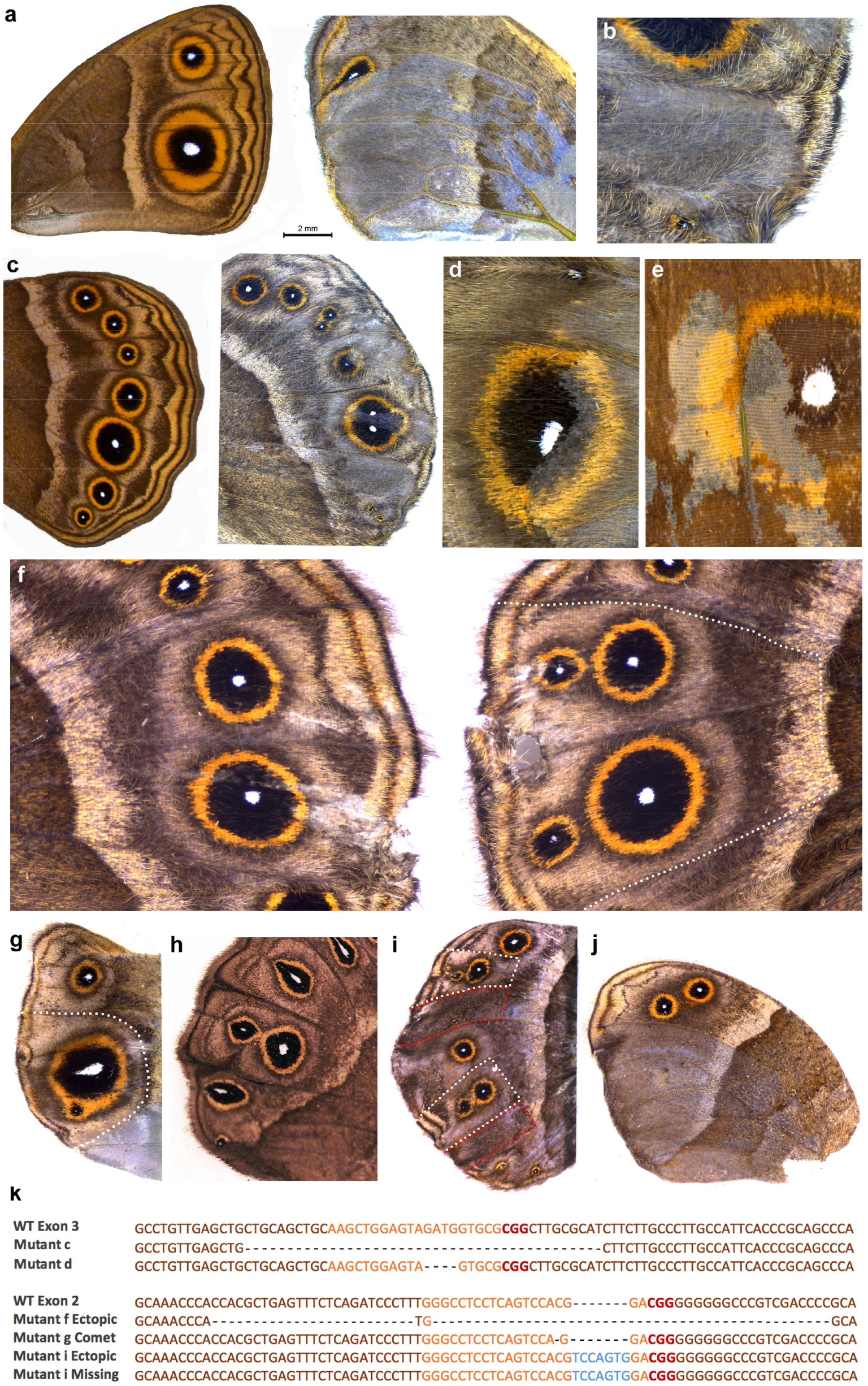
CRISPR mutants generated by targeting exon 2 and exon 3 of Dll. - **(a)** Wildtype forewing of *B. anynana* (left) Exon 3 phenotype with eyespots missing in areas of lighter pigmentation and disrupted venation (right). **(b)** Exon 3 phenotype with missing eyespot in a patch with mutant clones. **(c)** Wildtype hindwing of *B. anynana* (left) Exon 3 phenotype with split eyespots and bisected eyespot centers. **(d)** Exon 3 phenotype showing light colored scales in mutant clones across an eyespot. **(e)** Exon 3 phenotype with missing scales. **(f-j)** Exon 2 mutations. **(f)** Wildtype (left) and mutant wing (right) of the same individual where ectopic eyespots appeared on the distal hindwing margin after Exon 2 was targeted. **(g)** Comet shaped Cu1 eyespot center. **(h)** Example of a spontaneous comet mutant. **(i)** Wing with ectopic eyespots as well as missing eyespots. **(j)** Missing eyespots on hindwing in mosaic areas also showing lighter pigmentation. **(k)** Next generation sequencing of selected mutants (Exon 3 top panel and Exon 2 bottom panel) identifying the most frequent indels around the target site (Orange: guide region, Red: PAM sequence Blue: insertions, Dashed lines: deletions). Dotted lines on Exon 2 mutants in f, g and i represent wing regions carefully dissected for DNA isolation. Wing sectors for mutant i outlined in red (missing eyespots) were pooled for DNA isolation as were wing sectors outlined in white (ectopic eyespots). For Exon 3 mutant c the entire distal wing margin was dissected and for mutant d the area around the eyespot was dissected.

**Table 1:**
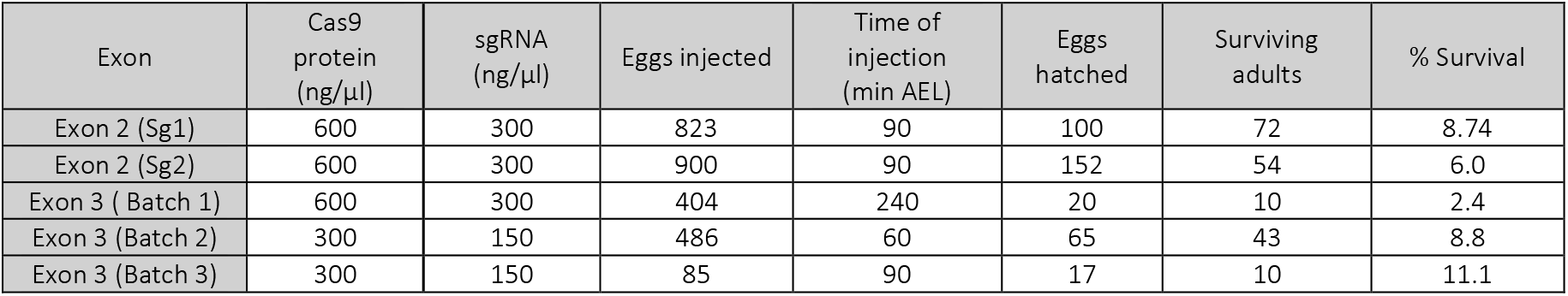
Embryo injection conditions and success rates for guide RNA injections targeting the two exons of *Dll*.

| Exon | Cas9 protein (ng/μl) | sgRNA (ng/μl) | Eggs injected | Time of injection (min AEL) | Eggs hatched | Surviving adults | % Survival |
| --- | --- | --- | --- | --- | --- | --- | --- |
| Exon 2 (Sg1) | 600 | 300 | 823 | 90 | 100 | 72 | 8.74 |
| Exon 2 (Sg2) | 600 | 300 | 900 | 90 | 152 | 54 | 6.0 |
| Exon 3 (Batch 1) | 600 | 300 | 404 | 240 | 20 | 10 | 2.4 |
| Exon 3 (Batch 2) | 300 | 150 | 486 | 60 | 65 | 43 | 8.8 |
| Exon 3 (Batch 3) | 300 | 150 | 85 | 90 | 17 | 10 | 11.1 |

**Table 2:** Overview of the mutant phenotypes observed in animals injected with *Dll* guide RNAs. Number of individuals displaying aberrations are reported. Total number of different eyespots carrying distortions are reported in brackets.

| Exon 2 | # Individuals examined | Incomplete eclosion | Margin disruption | Pigmentation disruptions | Split eyespots | Ectopic eyespots | Comet eyespots | Reduced eyespots | Missing eyespots | Leg/Antenna phenotypes |
| --- | --- | --- | --- | --- | --- | --- | --- | --- | --- | --- |
| Sg1 | 72 | 6 | 3 | 4 | 5 (5) | 3 (6) | 2(2) | 1 (1) | 3 (7) | 11 |
| Sg2 | 54 | 6 | 3 | 8 | 3 (2) | 3 (3) | 1(1) | 2 (2) | 3 (13) | 6 |

| Exon 3 | # Individuals examined | Incomplete eclosion | Margin disruption | Pigmentation disruptions | Split eyespots | Ectopic eyespots | Comet eyespots | Reduced eyespots | Missing eyespots | Leg/Antenna phenotypes |
| --- | --- | --- | --- | --- | --- | --- | --- | --- | --- | --- |
| Sg3 (1) | 10 | 0 | 0 | 0 | 0 | 0 | 0 | 0 | 0 | 1 |
| Sg3 (2) | 43 | 11 | 10 | 25 | 8 (12) | 0 | 0 | 0 | 2 (3) | 35 |
| Sg3 (3) | 10 | 3 | 4 | 6 | 3 (5) | 0 | 0 | 0 | 2 (3) | 6 |

* These results are based on easily visible phenotypes and are likely an underestimation particularly from individuals that partially or fully eclosed but with highly crumpled and folded wings making it difficult to evaluate the extent of the mutations.

### *Dll* exon 2 mutants produced gain and loss of function phenotypes

Embryonic injections of guide RNAs targeting either the 5′UTR (Sg1) or the coding sequence (Sg2) of exon 2 led to phenotypes similar to the ones described above (Table 2) as well as to a remarkable new set of phenotypes, sometimes co-occurring on the same wing. These included ectopic eyespots along the proximal-distal axis of the wing (Fig. 2f) and eyespots with a tear-drop shaped center (Fig. 2g), closely resembling a spontaneous mutant variant in *B. anynana* known as the comet phenotype^19^ (Fig. 2h). Ectopic eyespots were observed regardless of whether we targeted the 5′UTR or the coding sequence of exon 2, as we injected each of these guide RNAs separately. Some butterflies displayed both ectopic and missing eyespots on the same wing (Fig. 2i). Interestingly, ectopic eyespots were never associated with changes in pigmentation in contrast to wing tissue with missing eyespots (Fig. 2i,j), which always displayed the grey-blue pigmentation defects, highlighting the extent of the mutant clone of cells. Similarly to exon 3 mutants, we also observed appendage mutants including truncated antenna and legs or with fusion of antenna or proximal leg segments (Supplementary Fig. 2c,d,e).

### Confirmation of CRISPR-Ca9 activity using next generation sequencing

In order to confirm that the phenotypes observed were due to genetic alterations of the targeted exons, we performed next-generation amplicon sequencing of *Dll* to identify the entire range of mutations generated from Sg1, representing exon 2 mutations, and Sg3, representing exon 3 mutations. To identify mutations associated with each specific phenotype, especially in the case of exon 2 mutations that produced both ectopic as well as missing eyespots, we isolated DNA from the adult wing tissue by carefully dissecting around regions corresponding to missing, ectopic, or comet eyespots (see Fig 2f,g,i). To characterize mutations we used CRISPResso, a software pipeline for analyzing next generation sequencing data generated from CRISPR-Cas9 experiments^20^. This analysis identified a range of mutations from each wing tissue including deletions and insertions (Supplementary Figs 3, 4 and Supplementary Table 1); the most dominant mutations are shown in Figure 2k. The majority of mutations in exon 3 were comprised of frame-shift deletions whereas mutations induced by Sg1 were mostly non-coding (Table 2). For Sg3, we sequenced two individuals (Fig. 2c,d) and identified a range of mutations with the most frequent representing a 42 bp and 4 bp deletion, respectively (Fig. 2k and Supplementary Fig. 3a). For Sg1 we sequenced 3 individuals (Fig. 2f,g,i). A large 72bp deletion was observed in a mutant displaying ectopic eyespots (Fig. 2f, Supplementary Fig. 3b). In contrast, relatively small indels were observed for another ectopic eyespot mutant (Fig. 2i,k, Supplementary Fig. 3c), and surprisingly, the same 7 bp insertion emerged as the most dominant mutation from wing tissue either with ectopic or missing eyespots (Fig. 2i,k). The most dominant mutation observed for the comet eyespot phenotype represented a single base pair deletion (Fig. 2g,k, Supplementary Fig. 3b). Overall, CRISPResso identified only a very small proportion of mutations as disruptions to potential splice sites (0.1-0.2%). Because the link between specific mutations and the observed phenotypes was not clear, we decided to explore whether perhaps mutations that targeted each of the exons led to modifications in the way that *Dll* was transcribed.

### Targeting exon 2 induces alternative splicing

To explain the presence of ectopic eyespots following exon 2 CRISPR-Cas9 disruptions we examined the resulting cDNA sequences. RNA was isolated from embryos injected with each of the three guides (and Cas9) as well as from wild-type non-injected embryos. PCR amplification from cDNA using primers spanning exon 1 to exon 6 revealed that embryos injected with either Sg1 or Sg2, targeting exon 2, produced a novel product approximately 500 bp shorter than the wild-type product. Sequencing this short product revealed a deletion of 492 bp representing exon 2, suggesting that this exon had been completely spliced out. In contrast, we did not observe any alternative splicing for cDNA obtained from wild-type embryos or embryos injected with Sg3 (Supplementary Fig. 5).

To examine whether targeting exon 2 resulted in ectopic eyespots due to *Dll* overexpression we performed qPCR on cDNA from embryos injected either with Sg1 or Sg3, using primers designed to amplify exon 1. The aim of this experiment was to capture all *Dll* transcripts including the alternative spliced variants and to quantify them. The results did not reveal any significant differences in *Dll* expression after normalizing the data to our internal control gene EF1 alpha (Dll exon 1; p=0.66, EF1 alpha; p=0.08). Overall expression levels of *Dll* were low, with average Ct values of 29.88, S.E. = 0.43 (Sg) and 30.6, S.E. = 0.48 (Sg3) n = 4.

### Morphogens provide dynamic positional information in each developing wing sector

Several of the mutant *Dll* phenotypes suggested that this gene is involved in the process of eyespot center differentiation, which takes place during the late larval stage^12,21^ (Fig. 1c). Intriguing phenotypes involved the disappearance of eyespots, the splitting of the eyespot centers within a single wing sector bordered by veins, and the appearance of deformed eyespot centers and color rings near boundaries of wild-type and mutant tissue. In order to better understand how *Dll*, a transcription factor, might lead to such phenotypes we begun by examining the distribution of diffusible proteins (morphogens) able to provide complex spatial information within each wing sector. We tested whether two previously hypothesized morphogens, Wingless (Wg) and Decapentaplegic (Dpp), known to be involved in *Dll* regulation in early leg discs of *Drosophila^22^*, were expressed in *B. anynana* wing discs in the 5^th^ instar larvae. We cloned a 810 bp fragment for *B. anynana dpp* using specific primers (Supplementary Table 3) and performed *in situ* hybridizations. For visualization of Wg signaling, we looked at the expression of Armadillo (Arm) protein, a signal transducer of the Wg signaling pathway ^23^. In young 5^th^ instar wing discs we observed a *dpp* stripe in the middle of the wing discs, separating anterior from posterior wing compartments, as expected from work on *Drosophila* ^24^ (Supplementary Fig. 6a), but in slightly older larval discs, *dpp* was expressed across the whole wing, with slightly elevated expression in regions flanking each vein, and reduced expression in the future eyespot centers as well as in the midline of each wing sector (Fig. 3a, Supplementary Fig. 6b,c). At a late larval stage, *dpp* expression declines everywhere with the exception of the antero-posterior stripe (Supplementary Fig. 6d). Arm, on the other hand, was highly expressed in areas where *dpp* was missing, e.g., along the wing margin, and in the eyespot centers and in the midline (Fig. 3b, Supplementary Fig. 6e,f). We used information from these dynamic gene expression patterns, as well as from *Dll*, to model eyespot center differentiation.

**Figure 3.**
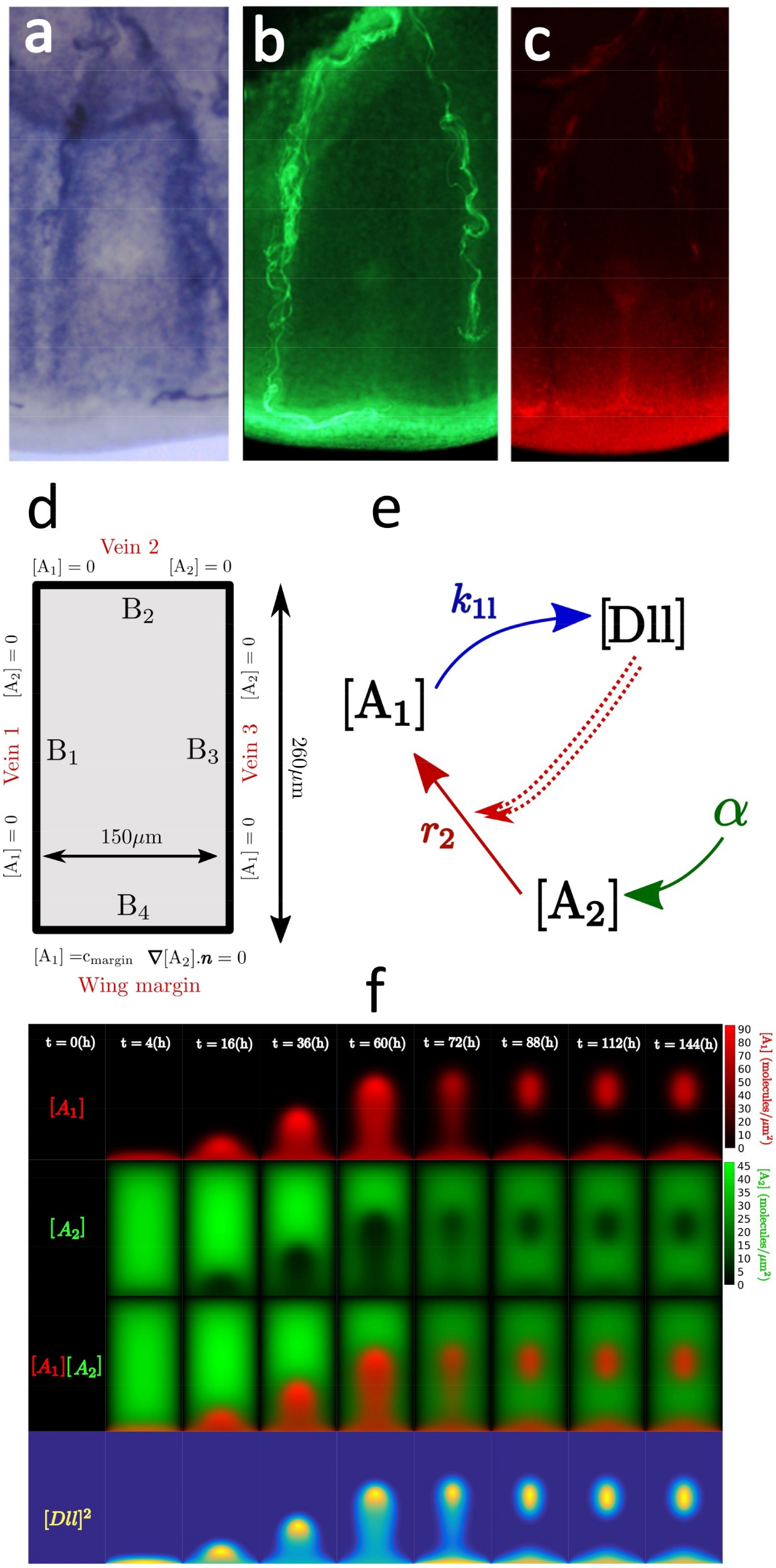
Morphogenetic inputs and modeling of eyespot formation. **(a)** *dpp* is expressed across the wing compartment but levels are lower in the eyespot centers. **(b)** Arm is located in eyespot centers, mid-line, as well as the wing margin. **(c)** *Dll* has a similar localization pattern to Arm. **(d)** Boundary conditions and size for the wing compartment. The boundaries with veins are modeled as sinks for both *A*_1_ and ***A***_2_. At the wing margin ***A***_1_ is imposed at a fixed concentration *c_margin_*, while we impose zero-flux conditions on ***A***_2_. **(e)** Interaction network involving the activator *A*_1_, the substrate *A*_2_ and *Dll. Dll* interacted cooperatively with itself (double dashed line) and with *A*_2_ to induce *A*_1_. During the reaction [A_2_]+2[Dll]->[A_1_], *A*_2_ was degraded. *A*_2_ is produced uniformly throughout the compartment. **(f)** Time-lapse results of reaction-diffusion simulation of eyespot centering. Concentration of A_1_ (first row) and A_2_ (second row) over 6 days. Third row is the overlap of A_1_ and A_2_. Fourth row represents square of Dll concentration, as this represents the Dll signaling incorporated within the model (see panel e).

### Gray-Scott model of eyespot formation

To probe the mechanism of eyespot center formation we utilized theoretical modeling to explore potential interactions between morphogens and the transcription factor *Dll*. In particular, our goal was to test whether such models could replicate the *Dll* mutant phenotypes by incorporating *Dll* as one of the components in the model and removing *Dll* function from patches of cells *in silico* that map to the patches of lighter coloration in *Dll* mutant wings.

We turned to a Gray-Scott reaction-diffusion model (also known as the Gierer-Meinhardt activator-depleted substrate model)^25,26^. A diffusible molecule A_1_ (putatively Wg) plays the role of an autocatalytic activator, and a diffusible molecule A_2_ (putatively Dpp) the role of a substrate that is degraded during activator production. A key difference between the Gray-Scott and activator-inhibitor reaction-diffusion models previously used for simulating spot formation - e.g. the Gierer-Meinhardt activator-inhibitor model,^16^ (Supplementary Theoretical Modeling) - is in how new eyespots form. In activator-inhibitor models, new activator maxima (i.e. eyespot centers) typically form between two existing maxima. However, in the Gray-Scott model, new maxima can form via a single maxima splitting ^27.^ The latter scenario appears to be closer to the experimental observations in *Dll* mutants as well as in spontaneous comet mutants of *B. anynana* (Fig. 2h, Supplementary Figs. 7,8,9). Another reason to not focus on activator-inhibitor type models is that our data suggested that *dpp* mRNA and Arm protein were anti-colocalized, which is counter to the assumptions underlying such models of eyespot formation ^26^.

### Incorporating *Dll* within the Gray-Scott model of eyespot formation

We included *Dll* within the network as a downstream gene activated by A_1_, which initially is expressed only along the wing margin. This is supported by the observation of co-localization of Arm and Dll (Fig. 3b,c) and by the assumption that the known activation of *Dll* by Wg (via Arm) in the *Drosophila* wing margin ^28^ is conserved in butterflies. A_2_ (Dpp) is uniformly produced throughout the wing compartment at a rate α, consistent with our *in situ* observations (Fig. 3b, Supplementary Fig. 6). *Dll* acts cooperatively with itself and conjointly with A_2_ to catalyze A_1_ production. These modeled interactions - though obviously a simplification - are compatible with data that show ectopic Dll activating endogenous *Dll* as well as *wg* in the wing and leg discs of *Drosophila^29^*. In this system, A_1_ production (e.g. Wg) is associated with A_2_ (e.g. Dpp) effective degradation, which could correspond to either real degradation or downregulation of A_2_ by A_1_ or by Dll. These interactions are summarized in Fig. 3e and they result in the anti-colocalization of A_1_ and A_2_, as experimentally observed. We emphasize that A_1_ and A_2_ do not correspond to single molecules, but more likely to sub-networks, with Wg and Dpp belonging to the sub-networks represented by A_1_ and A_2_ respectively.

The system we modeled is described by the following reaction-diffusion equations for the concentrations of *A*_1_ and *A*_2_, denoted by [*A*_1_ and [*A*_2_]. The action of Dll is included within the nonlinear reaction term (*K[A*_1_]^2^[*A*_2_]) (see Supplementary Theoretical Modeling for further details):

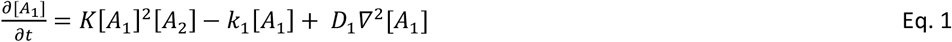

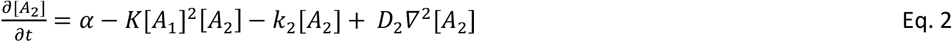

where *∇^2^* represents the two-dimensional Laplacian operator. The diffusion (*D*_1_ and *D*_2_), degradation (*k*_1_ and *k*_2_) and a (production of *A*_2_) rates are constrained by measurements in *Drosophila* ^30^ and *K* (representing the interaction between Dll, *A*_1_ and *A*_2_) is unknown. See Methods for details of boundary and initial conditions and Supplementary Theoretical Modeling for detailed description of simulation implementation and parameter tables. We constrained our parameter values to lie within the spot formation region of the phase space, where the reaction producing *A*_1_ degrades *A*_2_ at the same rate ^31,32^ Supplementary Fig. 8,10, Supplementary Theoretical Modeling).

## Gray-Scott model accurately replicates eyespot formation dynamics during larval stage

This reaction network produced a broad patch of activator (A_1_) up-regulation that narrows until it is along the midline and then further reduces to form a single spot, consistent with experimental observations (Fig. 3a,b,c,f)^33^. The eyespot location was near the observed experimental position using boundary conditions consistent with *in situ* observations (Fig. 3a,b,c,d). During the whole dynamics, A_1_ and A_2_ were spatially anti-correlated, in agreement with Arm and *dpp* anti-colocalization.

### Phase diagram of spot formation within the Gray-Scott model

The position, size, and shape of the spot within the model were sensitive to Dll activity (parameter K) and A_2_ production rate (parameter α) with eyespot centers emerging at high *K* and α (Supplementary Figs 7-8a). At lower values of *K* and α, the reaction between activator and substrate was not sufficiently strong to overcome degradation of the activator and no eyespot formed.

### Gray-Scott model accurately replicates eyespot formation of exon 3 mutant clones

We modeled *Dll* mutant clones as domains where *Dll* cannot be activated by A_1_. Mutant patches of *Dll* mutant cells were created within a simulated wing sector field by setting *K* to zero (red outlined regions in Fig. 4b). Outside this mutant patch, the reactions and boundary conditions remained unchanged. We assumed that A_1_ can diffuse within the *Dll* null region and that diffusion and production of A_2_ are not affected in that same region. We modeled seven *Dll* mutant clones where the mutant cells are present in different parts of the wing sector: (1) *Full*, covering the whole sector; (2) *Top*, covering the upper region of the wing compartment; (3) *Sliver*, present along one of the side veins; (4) *Diagonal*, distal from the wing margin; (5) *Comet*, distal from the wing margin but including part of the margin; (6) *Center*, present along a stripe at the center; and (7) *Corner*; present in two opposite corners of the sector (Fig. 4b,c,d,e,f,g,h). Our model was able to closely reproduce all the mutant phenotypes. For each phenotype we had the correct number of eyespot centers differentiated and they were positioned in close accord with our experimental observations. For comparison, we performed simulations of the same clones using the Gierer-Meinhardt activator-inhibitor type model, which reproduced most of the observed *Dll* mutant phenotypes, but did not have anti-colocalization of the two morphogens (A_1_ and A_2_) (Supplementary Fig. 13,14, Supplementary Theoretical Modeling).

**Figure 4.**
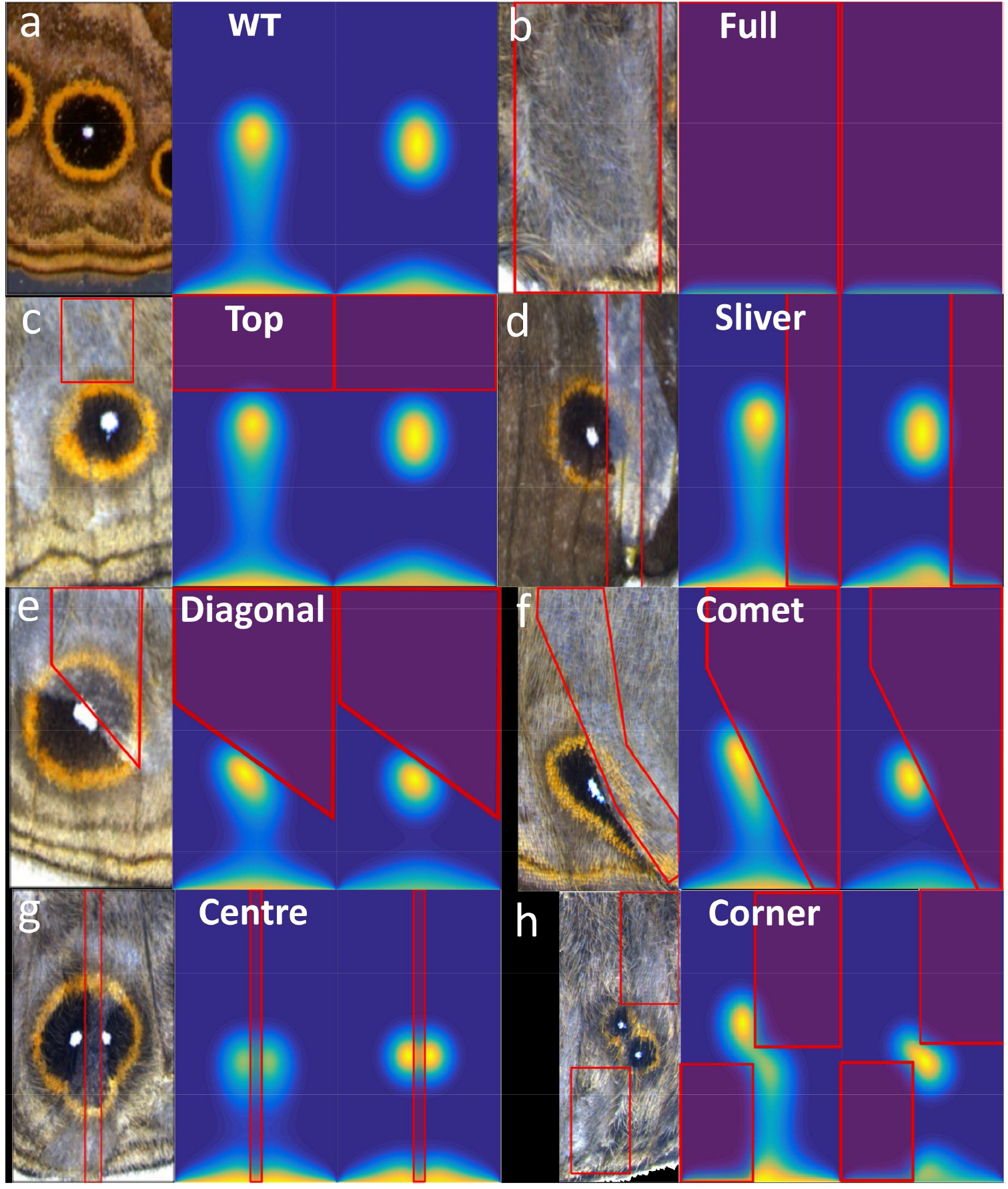
Reaction-diffusion simulations of wing sectors where part of the sector (red outline) has no "activator" function. For each panel, left image shows the experimental data. The right shows the *in silico* results after 72 hours and 144 hours, orientation of compartment and parameters are the same as in Fig. 3a. The region inside the red boxes in each images (except in a), represent the *Dll* mutant region, where *K=0*. See text for description of each phenotype.

### Our model accurately predicts ectopic eyespots and the comet phenotype from exon 2 mutants

Alternative splicing of exon 2 is associated with the differentiation of two eyespots and with comet-shaped eyespots. These phenotypes do not show associated pigmentation defects, and thus, it is unclear the extent of the *Dll* mutant clone that produced them. Therefore, we modeled these mutants by assuming cells expressed a functional truncated Dll protein across the whole wing sector, which degraded more slowly than its wild-type version, effectively resulting in increased *K* (Supplementary Theoretical Modeling). Increasing *K* while keeping all other parameters fixed led to a spot size increase, until at some threshold value the spot splits vertically into two smaller spots. This phenotype was very similar to the phenotypes observed in Fig. 2f,i, Fig.5b,c. Further increasing *K* resulted in the double spot phenotype turning into an extended finger pattern, close to the observed comet phenotype (Fig. 2g, Supplementary Figs. 8,9).

**Figure 5.**
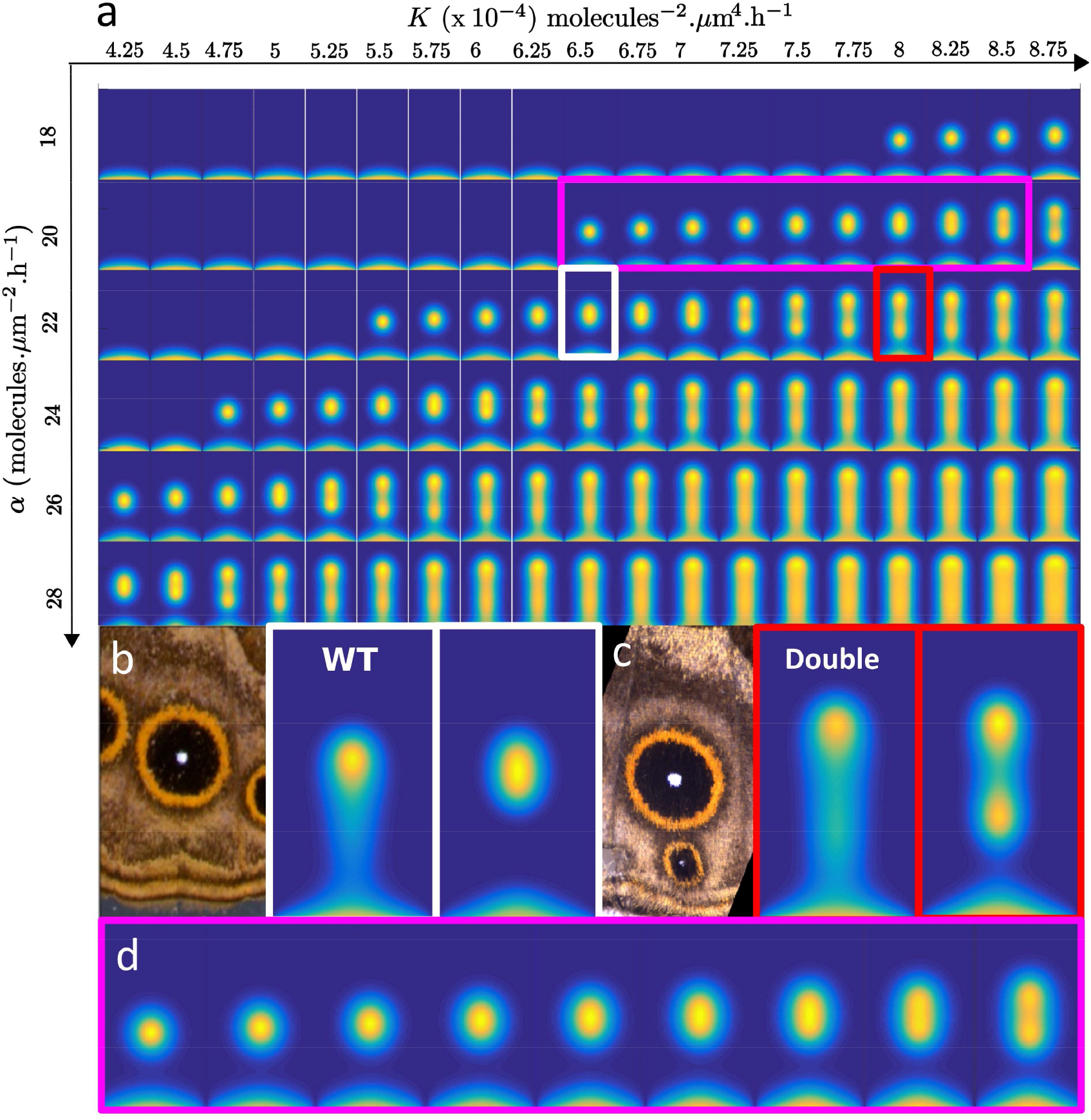
Perturbations of the Gray-Scott model and the K-a phase diagram reveal high sensitivity to changes in *Dll* functionality. **(a)** Phase diagram of A_1_ at t = 144h for different *K* and *α* parameters. **(b)** The parameters used for the wild-type correspond to the white rectangle in a. **(c)** Increasing *K* leads to appearance of a second spot. **(d)** Spot size increases when *K* increases with other parameters fixed, until spot splitting at high K. Region shown is close-up of pink region highlighted in panel a.

### Model predictions on eyespot size

Experimentally, it is known that reducing *Dll* expression across the whole wing results in reduced eyespot size^7^. Keeping our model parameters fixed, we reran the simulations with reduced values of K, which corresponds to reduced Dll production (Fig. 5a). The simulations support these experimental finding by showing that reducing *K* also results in smaller eyespots (Fig. 5d).

## Discussion

Gene expression studies have always showed a positive correlation between *Dll* expression and the number and size of eyespots that differentiate on the wings of different butterfly species, including *B. anynana* and *V. cardui* ^2^-^3^-^8^,^34^. During the larval stages, *Dll* is expressed in the center of the wing sectors where eyespots will develop, and is absent from the wing sectors where eyespots will not develop ^7,8^ In a recent study, Zhang and Reed (2016)^8^ found that CRISPR-Cas9 targeting exon 2 of *Dll* in *V. cardui* led to ectopic eyespots in wing sectors that normally display no eyespots, leading to the proposal that *Dll* must be a repressor of eyespot development. Mechanistically, however, this result is difficult to explain, as pointed out by the authors. Why would an eyespot repressor gene be naturally absent in sectors without eyespots and present in sectors with eyespots?

To explore this conundrum, we replicated these experiments in *B. anynana*. Similar to Zhang and Reed, we found that targeting the same regions of exon 2 resulted in butterflies with ectopic eyespots in addition to butterflies with missing eyespots, the latter of which was not observed in *V. cardui*. By exploring the effect of the guide RNAs on cDNA sequences obtained a few days after embryonic injections we found that disruptions in exon 2 produced transcripts completely lacking this exon, regardless of whether disruptions occurred in the 5′UTR or coding region of this exon. In contrast, several indels, but no exon skipping, occurred when targeting exon 3, indicating these disruptions led to a non-functional product. Furthermore, we only observed ectopic eyespots when targeting exon 2, suggesting that development of novel eyespots was a consequence of this exon splicing event.

Our simulation results predict that eyespot duplications can occur if the rate of Dll degradation is reduced across the wing cell, essentially leading to Dll overexpression. This led us to speculate that expression levels of *Dll* would be higher in embryos injected with Sg1 relative to Sg3. Our findings however did not support this hypothesis, possibly due to the overall low expression levels of *Dll* during embryonic development. Alternatively, it is possible that splicing is not impacting gene expression levels but is instead altering downstream processes affecting translation efficiency ^35^. The loss of exon 2 would have resulted in a shorter 5′UTR region, an alteration that is known to increase translation activity due to removal of inhibitory secondary structures and translational repressive elements ^36,37^.

A number of recent studies have shown that CRISPR-induced mutations can lead to alternative splicing and even gain of function phenotypes ^38-40^. Our results suggest that CRISPR-Cas9 targeting of exon 2 led to a truncated but potentially functional *Dll* transcript utilizing one of the numerous start codons present in exon 3 to produce an intact reading frame (Supplementary Fig. 5). In addition to ectopic eyespots, we also observed comet shaped eyespots which, in the spontaneous comet mutant, are associated with strong expression of *Dll*, suggesting an overexpression phenotype (Supplementary Fig. 9). Our modeling work also predicts comet phenotypes due to the emergence of a stable Dll finger as a result of increased protein expression. Future experiments will need to be performed for a fuller understanding of this phenomenon. In the meantime, it is interesting to note that a *Dll* over-expression phenotype may be easily achieved via disruptions to exon 2 of *Dll*.

A particularly curious observation was the emergence of butterflies with both missing and ectopic eyespots in different parts of the wing. By isolating wing tissue from these specific regions we hoped to correlate the frequency of a particular type of mutation with a particular phenotype (ectopic versus absent eyespots) using next-generation sequencing but we did not entirely succeed. Similar mutations were associated with both phenotypes. Our analysis also revealed that mutations associated with splice sites were extremely rare, thus exon skipping is likely occurring as a consequence of random DNA damage within this exon. It is well documented that exon skipping can be induced by a variety of mutations (nonsense, missense and silent) as well as varying sizes from single point mutations to large genomic deletions ^40^. Based on our findings, we propose that different phenotypes observed in adult wings may be related to the spatial distribution of each mutant cell clone in the wing sector, which cannot be inferred from the adult wing tissue and from particular mutation events inducing exon skipping, which so far we were unable to identify.

In contrast, guide RNAs targeting exon 3 led to typical missense mutations and to missing eyespots indicating that *Dll* is required for eyespot differentiation. A previous study performed in *B. anynana* had already functionally implicated *Dll* as a positive regulator of eyespot development, but the results were less stark than those reported here. *Dll* down-regulated via transgenic RNAi led to smaller eyespots, rather than missing eyespots, whereas its up-regulation led to two smaller eyespots appearing on the forewing ^7^. The *Dll* down-regulation failed to remove eyespots presumably because it was implemented during a limited period during late larval development and because it likely failed to eliminate *Dll* transcripts altogether - compared to CRISPR which can induce complete Dll-null clones. The two studies, however, by obtaining essentially the same results via the use of two different approaches, confirm that *Dll* is a positive regulator of eyespot development in *B. anynana* and likely also in other species.

Reaction-diffusion models have been shown to correctly recreate the patterning of vertebrate skin colors, of digits in mice, and of distal fin elements in catsharks^41-43^. Our reaction-diffusion simulations - and in particular our ability to simulate the mutant phenotypes by simply implementing a region where the reaction term for Dll is removed - enabled us to dissect the processes by which eyespots emerge. Our modeling also supports a likely functional role for the morphogenetic ligands Wg and Dpp in eyespot formation, size, and positioning. An important corollary of our results is that activator-inhibitor models may not be suitable for studying eyespot patterning. The anti-localization of the Wg and Dpp signals suggests that activator-substrate models - such as the Gray-Scott model - may be more applicable to modeling eyespot patterning.

In addition to eyespot center differentiation, we confirmed that *Dll* has an additional role in wing melanization, previously shown in *B. anynana* ^7^, as well as in *Drosophila biarmipes* and *Junonia orythia* ^9, 44^. In *B. anynana*, ectopic expression of *Dll* during the early pupal wing led to patches of darker scales on the wing, whereas *Dll* RNAi led to no observable change in color ^7^. The current *Dll* exon 3 mutants, with light colored patches of pigmentation, lend additional critical support for this function of *Dll*.

*Dll* appears to have a further role in scale cell development. In several *Dll* mutants, a specific type of scale, the cover scales, or both cover and ground scales were missing from patches on the wing. Patches of scales with reduced pigmentation may have been *Dll* heterozygous clones, whereas those with scales missing may have been homozygous clones. This suggests that *Dll* is required for scale development. Scale cells, due to their pattern of division, differentiation and growth, and expression of an *achaete-scute* homologue, were proposed to be homologous to *Drosophila* sensory bristles, which share similar characteristics but are restricted to the anterior margin in the fly wing ^45^. In *Drosophila, Dll* mutant clones along the wing margin lead to loss of *achaete-scute* expression and loss of bristles ^46^. Our results further strengthen the hypothesis that butterfly wing scales are novel traits that originated from modified sensory bristles, which populated the entire wing blade.

## Conclusions

Here, we show that CRISPR cas-9 induced mutations in *Dll* can produce both knockout and gain of function phenotypes depending on which specific exon is targeted. While we still do not understand which mutations lead to the different mutant phenotypes, we propose that gain of function phenotypes are associated with alternative splicing of *Dll*. Our results also demonstrate that *Dll* is required for eyespot differentiation and for the production of melanin pigmentation across the whole wing, not just in the black regions of the eyespot, where its expression is stronger. Furthermore, our work confirms the involvement of *Dll* in ventral appendage development in *B. anynana*, and additionally in scale development, a function not previously reported for this gene. Finally, we provide a detailed reaction-diffusion model that accurately describes the dynamics of both wild-type and mutant eyespot formation, and identify the first molecule, *Dll*, playing a role in this reaction-diffusion process.

## Acknowledgements

HC and JVC were funded by the Ministry of Singapore grant MOE2014-T2-1-146 and MOE2015-T2-2-159 awarded to AM; ST was funded by a HFSP Young Investigator Grant (RGY0083/2016) awarded to TES. TES was also supported by a National Research Foundation Fellowship (NRF2012NRF-NRFF001-094). We gratefully acknowledge Kathy Su, Gowri Rajaratnam and Rudolf Meier (DBS, NUS) for facilitating Next Generation sequencing and for stimulating discussions on the research. We also thank Nick Tolwinsky for providing the Arm antibody, Yuji Matsuoka for technical advice on CRISPR experiments and Firefly farms, Singapore, for the corn supply.

## Author Contributions

H. C., S.T, T.E.S and A.M designed the study and wrote the paper. S.T, T.Y.J.L: reaction-diffusion modeling. H.C: CRISPR/Cas9 genome editing on exon 2, cDNA synthesis, gene cloning, qPCR and Next Generation Sequencing analysis. J.vC: CRISPR/Cas9 genome editing on exon 3. T.B: *In situ* hybridization and antibody stainings. All authors read and approved the final manuscript.

## Materials and Methods

### Animal husbandry

*B. anynana* were reared at 27^o^C and 60% humidity inside a climate room with 12:12hrs light: dark cycle. All larvae were fed young corn leaves until pupation. Emerged butterflies were frozen and then the wings were cut from the body for imaging using a Leica DMS1000 digital microscope.

### Guide RNA design

Guide RNAs corresponding to GGN_20_NGG (Dll) were designed using CRISPR Direct ^47^. We separately targeted three sites in Dll with two guides targeting exon 2, (in the 5′UTR and coding sequence) and a third guide targeting the homeobox of exon 3 (Fig. 1a). The guide RNAs were created by amplifying overlapping primers ^48^ (Supplementary Table 3) using Q5 polymerase (New England Biolabs). One primer contains the T7 promoter sequence and gene target region and the other is a common reverse primer composed of the guide RNA backbone. Constructs were transcribed using T7 polymerase and (10X) transcription buffer (New England Biolabs), RNAse inhibitor (Ribolock), NTPs (10mM) and 300ng of the guide template. Final sample volume was 20 µl. Samples were incubated for 16h at 37°C and then subject to DNase treatment at 37°C for 1 hour. Samples were purified by ethanol precipitation and RNA size and integrity was confirmed by gel electrophoresis.

### *In vitro* cleavage assay

The guide RNAs were tested using an in vitro cleavage assay. Wildtype genomic DNA was amplified using primers designed to the region flanking the guide RNA target sites. Guide RNA (160ng) and Cas9 protein (322ng), 10X buffer (1 µl) were brought to a final volume of 10 µl with nuclease free water and incubated for 15 mins at 37°C. The purified amplicon (100ng) was added and the reaction incubated for a further 1-2h at 37°C. The entire reaction volume was analysed on a 2% agarose gel. Cas9 protein was purchased from two suppliers, NEB EnGen Cas9 NLS (Exon 2 injections) and PNA Bio Inc. (Exon 3 injections).

### Embryo injections

Wildtype *B. anynana* adults were allowed to lay eggs on corn plants. Eggs were picked within one hour after oviposition and immobilised with 1mm wide strips of double sided tape in plastic 90mm petri dishes. Cas9 protein and guide RNA were prepared in a 10µl volume and incubated for 15 mins at 37°C prior to injection along with 0.5 µl of food dye to aid embryo injections (Table 1). The mixture was injected into eggs by nitrogen driven injections through glass capillary needles. Injected eggs were stored in closed petri dishes, accompanied by daily re-dampened cotton balls to maintain humidity. After hatching, larvae were reared in small containers for one week then moved to corn plants to complete their development.

### Screening and genotyping mutants

Upon emergence, butterflies were immediately stored at-80°C in individual containers. All individuals were screened under a microscope and examined for asymmetric mutant phenotypes. For selected mutants, genomic DNA was extracted from dissected wing tissue displaying mutant clone regions and modified/ectopic eyespots (E.Z.N.A tissue DNA kit). For next generation sequencing, amplicons shorter than 500 bp incorporating exon 2 or exon 3 were amplified using barcoded primers by PCR (Supplementary Table 3). The samples were visualized on a gel to confirm the presence of a single band then purified using Thermo Scientific PCR purification kit. The purified products were quantified using Qubit and sequenced using Illumina Miseq (300 bp paired-end). Exon 3 mutants were sequenced with AIT (Singapore), and exon 2 mutants were sent to GIS (Singapore). Sequencing coverage was 10,000x. Demultiplexing was performed on an in-house python script ^49^. The fastq files were checked for quality and trimmed using PRINSEQ ^50^. The trimmed files were processed using the command line version of CRISPResso ^20^.

### Detection of alternative splicing and quantitative PCR

RNA was isolated from injected eggs (guides targeting 5′UTR of exon 2 (Sg1) and the homebox domain of exon 3 (Sg3)) and control eggs (no injection) using Qiagen RNeasy mini kit incorporating a DNase 1 treatment (Thermo Fisher Scientific). RNA was isolated 2 days after egg injection. For each treatment group, we prepared four replicates of 50 pooled eggs on the same day. To control for developmental timing, we alternated injecting 50 eggs between the two groups (Sg1 and Sg3) for a total of 200 eggs/group. Eggs were placed in a petri dish of PBS and injected within 90 minutes of oviposition. After 2 days, eggs were carefully removed from the PBS and briefly transferred to RNAlater^®^ (Qiagen) prior to RNA isolation. For each of the 12 RNA samples, 2µg of RNA was used as input for cDNA synthesis (Thermo Scientific Revertaid First Strand). cDNA was also obtained from 50 pooled eggs injected with Sg2 in a separate experiment using the same protocol. A PCR was performed on the cDNA using *Dll* primers spanning exons 1-6 (wild-type = 1.5kb product) and visualized on a 1.5% agarose gel. The spliced transcript produced from the guide targeting the 5′UTR of exon was cloned into a pGEM t-easy vector followed by colony PCR using M13 primers to identify colonies carrying this product. The short insert (~1kb) was amplified using Big Dye sequencing kit (Thermo Fisher Scientific) and sequenced.

qPCR was performed on the cDNA from embryo’s injected with Sg1 or Sg3 (representing exon 2 and exon 3 disruptions). Primers were designed using Primer3 plus for *Dll* exon 1 and an internal control gene EF1 alpha. Relative expression was performed using a qPCR mastermix (Kapa SYBR Fast Uni) and 4ng of cDNA from 4 biological replicates and 2 technical replicates in a single experiment. Four biological replicates were tested to ensure sufficient statistical power to detect expression differences. The reaction was set up following the manufacturer’s instructions and run on a BIORAD thermocycler. Relative expression software tool (REST) was used to analyze the expression data ^51^.

### In-situ Hybridization

In-situ hybridization was performed on 5^th^ instar larval wing discs. Wings were dissected in cold PBS and transferred into fixative containing 4% formaldehyde. After proteinase K treatment peripodial membranes were removed using fine forceps. The wings were then gradually transferred in increasing concentration of pre-hybridization buffer in PBST and incubated in pre-hybridization buffer at 65°C for 1 hr before transferring into hybridization buffer containing 70ng/ml probe. Hybridization was carried out in a rocking-heating incubator at 65°C for 20 hrs. After hybridization wings were washed 5 times in pre-hybridization buffer for 20 mins at 65°C. Blocking was carried out using 1% BSA in PBST. Anti-digoxygenin AP(Roche) at the concentration 1:3000 was used to tag digoxygenin labelled probes. NBT/BCIP (Promega) in alkaline phosphatase buffer was used to generate color. Imaging was carried out under Leica DMS1000 microscope using LAS v4.9 software.

### Antibody staining

5^th^ instar larval wing disc were dissected in cold PBS and incubated in fix buffer containing 4% formaldehyde for 35 mins, washed 4 times in cold PBS and blocked using block buffer for 2 days. Wings were stained against Armadillo using primary antibody (294 rabbit anti-Arm; a gift from Nicholas Tolwinski ^52^) at the concentration of 1:10 and secondary antibody (alexa fluor 488 goat anti-rabbit: Thermo) at the concentration of 1:800. Wings were then mounted on prolong gold antifade reagent (Thermo) and imaged under Zeiss Axio Imager M2 using Zen 2012 software.

### Modeling details

#### Parameter estimation

We modeled a wing sector bordered by veins and containing a single eyespot as a rectangle with typical width *L_x_ =* 150 *µm* and length *L_y_ = 262 µm* (Fig. 3d),^16^. We used degradation and diffusion rates for both A_1_ and A_2_ close in magnitude to those measured for Wg and Dpp respectively in the *Drosophila* wing disc ^30^. Due to the longer time scales involved in eyespot patterning, both degradation and diffusion rates were assumed to be smaller than in *Drosophila* (therefore, we explored values varying by a factor of 0.1 to 1). In line with experimental observations where we observed a decrease in *dpp* (Supplementary Fig. 6) at late larval stage, we decreased α by 25% at time t = 60h in the simulation.

We present in Fig. 4 the results of the simulations for the different *Dll* mutant conditions. Results are shown for the parameter set that maximizes the matching between eyespot number and location(s) in the wing compartment between the simulations and the experimental data. The same parameter set was used in all simulation results shown. To model Exon 2 mutations, we increased K, which corresponds to either increasing *Dll* expression levels or decreasing its degradation rate (See Supplementary Theoretical Modeling).

#### Boundary conditions

Boundary conditions were implemented based on the *in situs* and immunostainings for *dpp* and Arm (Fig. 3a,b). The wing margin was modeled as a source term of *Wg* as Arm is present along the wing margin of *B. anynana* and *wg* is also present along the wing margin of other butterflies ^53^. As *dpp* is absent along the wing veins (Fig. 3a), we modeled the veins as sinks for both Wg and Dpp, which helped to confine the activator and substrate to the central part of the wing sector in a finger-like pattern (Fig.3d,f). These conditions differ from those used in ^15,16^ where the proximal cross-vein and lateral veins are the only sources of activator and inhibitor.

#### Initial conditions

At t = 0h, there are no activator and substrate in the wing sector. At t = 0h, A_1_ starts to diffuse from the wing margin to the wing sector, and the substrate A_2_ is produced by all cells in the wing sector. We assume detailed balance in the reactions, which can lead to spot formation in the Gray-Scott model (Supplementary Theoretical Modeling).

## Data availability

The codes used to generate simulations and the images of all wings from butterflies showing phenotypes are available from the corresponding authors upon request.

